# BDNF-dependent modulation of axonal transport is selectively impaired in ALS

**DOI:** 10.1101/2021.12.06.471484

**Authors:** Andrew P. Tosolini, James N. Sleigh, Sunaina Surana, Elena R. Rhymes, Stephen D. Cahalan, Giampietro Schiavo

## Abstract

Axonal transport ensures long-range delivery of essential cargoes between proximal and distal compartments of neurons, and is needed for neuronal development, function, and survival. Deficits in axonal transport have been detected at pre-symptomatic stages in mouse models of amyotrophic lateral sclerosis (ALS), suggesting that impairments are fundamental for disease pathogenesis. However, the precise mechanisms responsible for the transport deficits and whether they preferentially affect α-motor neuron (MN) subtypes remain unresolved. Here, we report that stimulation of wild-type neurons with brain-derived neurotrophic factor (BDNF) enhances trafficking of signalling endosomes specifically in fast MNs (FMNs). In early symptomatic SOD1^G93A^ mice, FMNs display selective impairment of axonal transport and develop an insensitivity to BDNF stimulation, with pathology upregulating classical non-pro-survival receptors in muscles and sciatic nerves. Altogether, these data indicate that cell- and non-cell autonomous BDNF signalling is impaired in vulnerable SOD1^G93A^ MNs, thus identifying a new key deficit in ALS amenable for future therapeutic interventions.

## Introduction

Amyotrophic lateral sclerosis (ALS) is a progressive neurodegenerative disease primarily affecting motor neurons (MNs), leading to muscle atrophy, paralysis and ultimately death due to respiratory failure. Although only a small proportion of ALS-causing mutations are found in genes encoding axonal transport components (e.g., *KIF5A, DCTN1, ANXA11*), altered axonal transport is a common pathological feature downstream of many ALS-causing mutations (Brown and Al-Chalabi, 2017; De Vos and Hafezparast, 2017). Axonal transport maintains neuronal homeostasis by ensuring the long-range delivery of several cargoes, including cytoskeletal components, organelles, signalling molecules and RNA between proximal and distal neuronal compartments (Terenzio et al., 2017). As a result, perturbations in axonal transport have severe consequences for neuronal homeostasis and function (Sleigh et al., 2019).

α-MNs are defined by the type of skeletal muscle fibre they innervate, and can be sub-classified according to their firing pattern into fast (FMNs) and slow (SMNs) MNs, each with distinct anatomical, metabolic, and functional properties (Kanning et al., 2010; Stifani, 2014; Ragagnin et al., 2019), as well as diverse transcriptional profiles (Blum et al., 2021). FMNs innervate type-IIb and -IIx fast-twitch fatigable and type IIa fast-twitch fatigue-resistant muscle fibres to execute fine motor control, whereas SMNs innervate type I slow-twitch fatigue-resistant muscle fibres to exert postural control (Stifani, 2014). Strikingly, FMNs are more susceptible to ALS pathology, whereas SMNs are predominantly resistant (Nijssen et al., 2017). This FMN preferential vulnerability has been observed in SOD1 (Frey et al., 2000; Pun et al., 2006; Tremblay et al., 2017; Allodi et al., 2021), TDP-43 (Ebstein et al., 2019), FUS (Sharma et al., 2016) and C9ORF72 (Liu et al., 2016) mutant mice, with limb-onset ALS accounting for ∼70% of human pathology (Kiernan et al., 2011), suggestive of preferential FMN vulnerability conserved across species.

The tibialis anterior muscle (TA), exclusively innervated by FMNs (Delezie et al., 2019), is affected early in ALS mice, with neuromuscular junction (NMJ) denervation occurring before MN loss (Alhindi et al., 2021), and is similarly impacted in ALS patients (de Carvalho and Swash, 2013; Jenkins et al., 2020). In contrast, the predominantly SMN-innervated soleus muscle is more resistant to pathology (Alhindi et al., 2021). ALS pathology also induces a fast-to-slow muscle fibre type switch (Hegedus et al., 2008), a phenotype that is similar to mice lacking muscle-specific brain-derived neurotrophic factor (BDNF) (Delezie et al., 2019).

The neurotrophin BDNF controls the development and maintenance of neurons through binding to TrkB and p75^NTR^ receptors. Full-length TrkB (TrkB.FL) contains a cytoplasmic tyrosine kinase domain essential for pro-survival signalling via ERK1/2, Akt and PLC-*γ* controlled pathways (Chao, 2003). Activation of these pathways is dampened by a truncated TrkB receptor isoform (TrkB.T1) lacking the kinase domain, which sequesters synaptic BDNF (Haapasalo et al., 2002). The physiological roles of p75^NTR^ are equally complex, with high affinity for pro-neurotrophins that control neuronal apoptosis during development, whilst modulating neurotransmitter availability and NMJ organisation in the mature nervous system (Pérez et al., 2019). BDNF binding triggers TrkB.FL, TrkB.T1 and p75^NTR^ homo- and/or hetero-dimerisation (Skeldal et al., 2011), and each complex elicits distinct signalling outputs (e.g., TrkB.FL-TrkB.T1 heterodimers inhibit TrkB.FL autophosphorylation (Hurtado et al., 2017)).

Despite in-depth knowledge of BDNF biology (Chao, 2003), the physiological landscape of BDNF signalling at the NMJ, as well as its possible perturbation in ALS, are currently unknown. BDNF is secreted by skeletal muscle during contraction (Gómez-Pinilla et al., 2002; Hurtado et al., 2017), and regulates both the pre- and post-synaptic components of the neuromuscular synapse (Delezie et al., 2019). Internalised BDNF-receptor complexes induce local signalling (Chao, 2003) and local translation (Santos et al., 2010), and are sorted to signalling endosomes, which undergo fast retrograde transport to the soma (Deinhardt et al., 2006), with signalling endosome flux dependent on TrkB activation (Wang et al., 2016). Hence, understanding the regulation of BDNF-signalling in MN subtypes can provide novel clues regarding selective MN vulnerability in ALS.

Here, we assessed axonal transport dynamics of signalling endosomes in axons of different α-MN subtypes in WT and ALS mice *in vivo*. We find that WT FMNs display faster retrograde transport of signalling endosomes when stimulated with BDNF. In SOD1^G93A^ mice, transport is preferentially impaired in FMNs innervating TA, and develop an insensitivity to BDNF stimulation. In addition, we show that TrkB.T1 and p75^NTR^ expression is upregulated in SOD1^G93A^ muscles, sciatic nerves and Schwann cells thus identifying cell- and non-cell-autonomous dysregulation of BDNF signalling in ALS pathology.

## Results

### MN subtypes display differential regulation of axonal transport by BDNF

To investigate the influence of α-MN and skeletal muscle subtypes (Kanning et al., 2010; Stifani, 2014; Ragagnin et al., 2019) on axonal transport dynamics, we labelled neurotrophin-containing signalling endosomes with a fluorescent atoxic tetanus neurotoxin binding fragment (HcT) (Surana et al., 2018, 2020) to assess axonal transport dynamics in sciatic nerves of live mice (Sleigh et al., 2020a). H_C_T is internalised in MNs upon binding to nidogens and polysialogangliosides (Bercsenyi et al., 2014), and is retrogradely transported in Rab7-positive signalling endosomes (Deinhardt et al., 2006). Using H_C_T-555, we separately targeted TA, lateral gastrocnemius (LG) and soleus muscles in wild-type (WT) mice, and after 4-8 hours, we performed time-lapse microscopy (**Figure 1A**). Speed distribution curves (**Figure 1B**) and the mean and maximum velocities of signalling endosomes (**Figure 1C; Figures S1A-B**) indicate that the transport dynamics in axons supplying TA, LG and soleus are similar across healthy neurons. However, less pausing was observed in TA axons (**Figure 1D**). Using previously reported methods (Sleigh et al., 2020b), we show that the mean diameter of motor axons innervating the TA, LG and soleus are similar (**Figure 1E; Figure S1C**).

**Figure 1.**
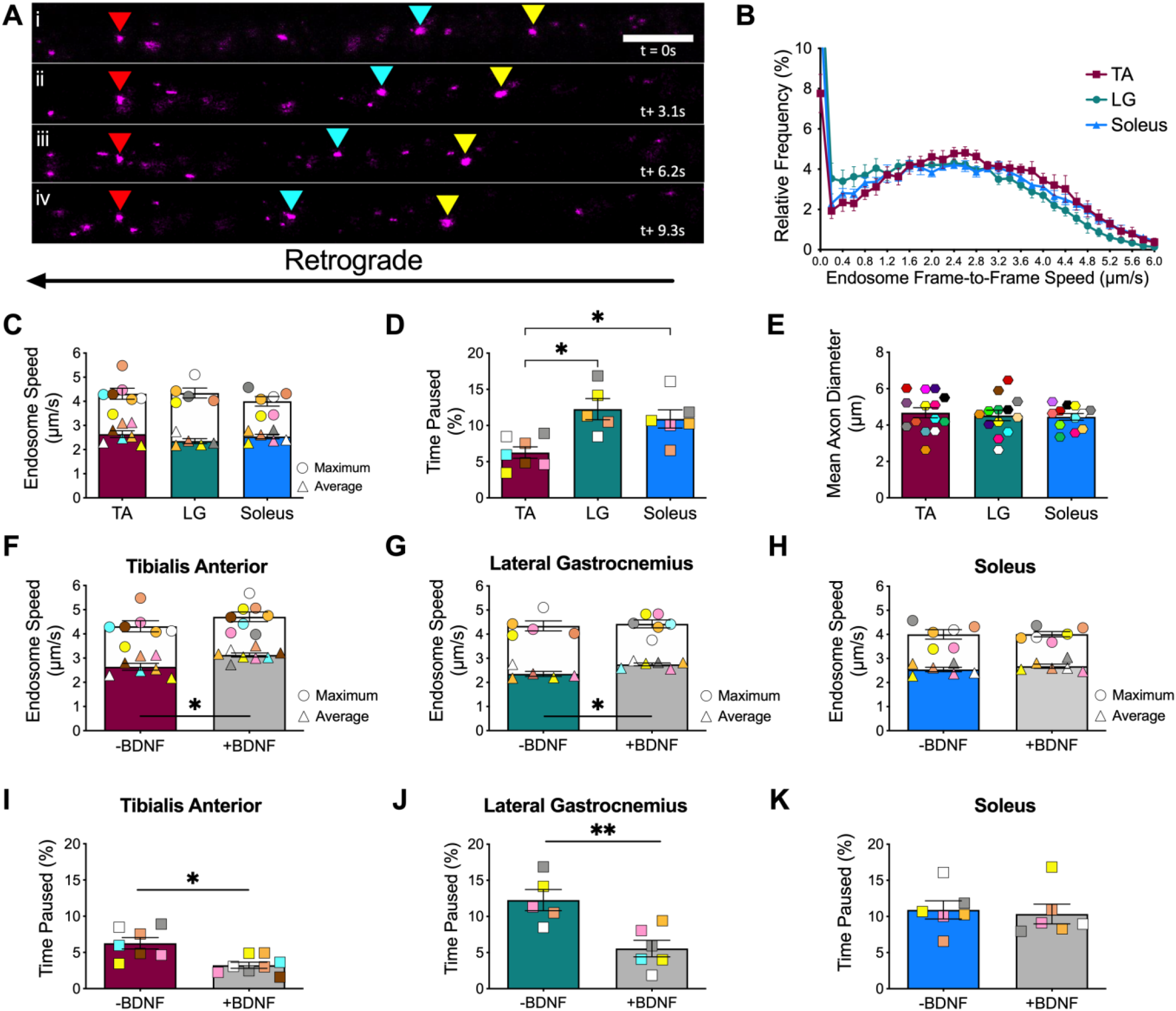
BDNF stimulation differentially impacts axonal transport of signalling endosomes in motor neurons innervating fast and slow muscles in wild-type mice. **A)** Time-lapse images of retrogradely transported (i.e., right-to-left), H_C_T-555-containing signalling endosomes (magenta) from a single sciatic nerve axon. Yellow and cyan arrows indicate retrogradely moving signalling endosomes, while red arrows identify a paused endosome. Frame rate = 3.1 frame/s; scale bar = 10 μm. **B)** Endosome frame-to-frame speed distribution curves in motor axons innervating tibialis anterior (TA), lateral gastrocnemius (LG) and soleus muscles in basal conditions. **C)** Mean (triangles) and maximum (circles) endosome speeds in motor axons innervating TA, LG, and soleus (p=0.193, one-way ANOVA, n=5-7). **D)** Relative percentage of time signalling endosomes paused in axons innervating the TA, LG, and soleus (p=0.002, one-way ANOVA, n=5-7). **E)** Mean diameter of H_C_T-containing motor axons innervating TA, LG, and soleus (p=0.819, one-way ANOVA, n=16-21). **F)** Mean and maximum endosome speeds upon intramuscular BDNF stimulation in motor axons innervating TA (p=0.029), **G)** LG (p=0.017), and **H)** soleus (p=0.485). **I)** Time signalling endosomes paused upon BDNF stimulation in motor axons innervating TA (p=0.014), **J)** LG (p=0.009) and **K)** soleus (p=0.559). *F-K)* were assessed by a Mann-Whitney test (n=5-8). Means ± SEM are plotted for all graphs. Colour coding is consistent and reflects the same animal in **C-D** and **F-K**. *p<0.05, **p<0.01. See also **Figures S1-S2**.

We next assessed whether stimulation with BDNF impacts signalling endosome transport dynamics, given BDNFs influence on endocytosis, endosomal flux and pro-survival signalling events (Wang et al., 2016; Bronfman and Moya-Alvarado, 2020). Co-injection of H_C_T-555 with 25 ng of BDNF increased the mean speeds of signalling endosomes in motor axons innervating TA (**Figure 1F; Figure S1D**) and LG (**Figure 1G; Figure S1E**), whilst concurrently reducing their pausing time (**Figures 1I-1J)**. However, BDNF application had no influence on transport in soleus motor axons (**Figures 1H, 1K; Figure S1F**). We then tested if this response was specific for BDNF or a general feature of neurotrophins, by stimulating FMN axons with glial cell line-derived neurotrophic factor (GDNF), which can activate distinct signalling cascades via RET and GFRα receptors (Airaksinen and Saarma, 2002). In contrast to BDNF, application of 25 ng of GDNF did not influence transport of H_C_T-555-positive signalling endosomes (**Figures S1G-H)**. Altogether, these data indicate that FMNs and SMNs have similar axonal transport dynamics under basal conditions, and that BDNF stimulation enhances axonal transport specifically in FMNs.

### Axonal transport is selectively impaired in TA-innervating axons of SOD1^G93A^ mice

SOD1^G93A^ mice display early and persistent axonal transport deficits (Bilsland et al., 2010), but the precise contributions of fast and slow MNs, as well as BDNF stimulation, are unresolved. First, we assessed *in vitro* axonal transport in mixed embryonic primary ventral horn cultures (**Figure S2Bi)**, and observed similar transport speeds between WT and SOD1^G93A^ neurons (**Figure S2Bii**), thus supporting a neurodegenerative, rather than neurodevelopmental, transport phenotype in SOD1^G93A^ mice (Bilsland et al., 2010). However, application of 50 ng/ml of BDNF increased retrograde transport speeds in WT, but not SOD1^G93A^, primary ventral horn neurons, suggestive of dysregulated BDNF signalling in SOD1^G93A^ mice (**Figures S2Biii-v**).

We then assessed *in vivo* axonal transport dynamics at postnatal day 73 (P73) and 94 (P94), which correspond to disease time-points characterised by ∼20% and ∼40% loss of lumbar MNs, respectively (Bilsland et al., 2010). Axonal transport of signalling endosomes was impaired in TA-innervating motor axons at both timepoints (**Figure 2A)**, and without significant alterations in pausing (**Figure S2Ci)**. On the other hand, axonal transport was unaffected in motor axons innervating the soleus (**Figure 2B; Figure S2Di)** or LG (**Figures S2Ei-iii**) in SOD1^G93A^ mice.

**Figure 2.**
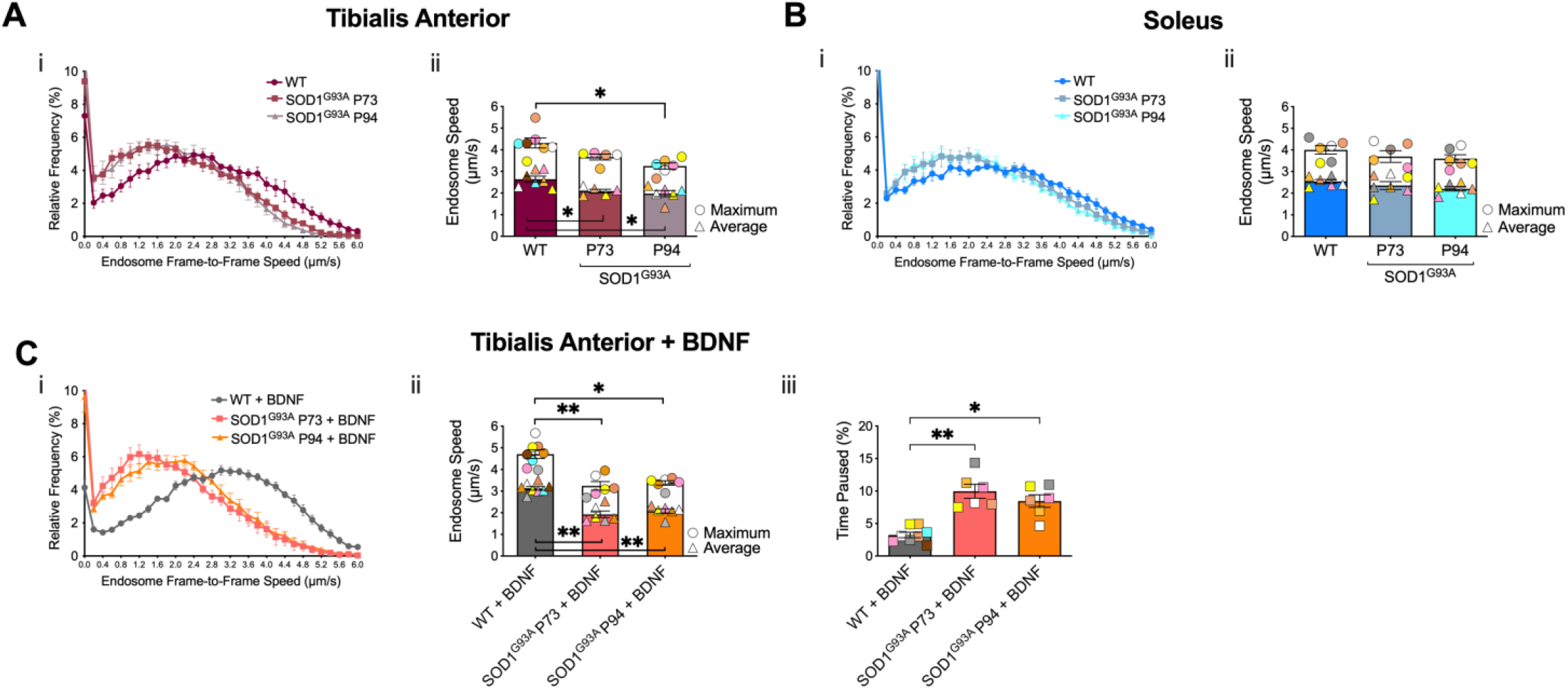
Retrograde transport and sensitivity to exogenous BDNF are selectively impaired in motor axons innervating TA in SO D1^G93A^ mice. Retrograde transport in motor axons innervating **A)** TA and **B)** soleus at two stages of SOD1^G93A^ disease (P73 and P94) compared to wild type (WT) mice, displaying the *i)* endosome frame-to-frame speed distribution curves, and *ii)* mean and maximum endosome speeds (TA: p=0.004; soleus: p=0.28, one-way ANOVA, n=5-7). **C)** Axonal endosome transport in P73 and P94 SOD1^G93A^ vs. WT mice upon intramuscular BDNF application in motor axons innervating TA, displaying *i)* endosome frame-to-frame speed distribution curves, *ii)* mean and maximum endosome speeds (p<0.0001, one-way ANOVA), and *iii)* relative percentage of time signalling endosomes paused (p=0.0001). Means ± SEM are plotted for all graphs. *p<0.05, **p<0.01, Kruskal-Wallis multiple comparisons test (n=6-8). See also **Figure S2**.

We then evaluated the impact of BDNF stimulation on transport in SOD1^G93A^ mice. In contrast to WT mice, BDNF stimulation failed to enhance transport speeds in motor axons innervating TA (**Figures S2Cii-iii**) or LG (**Figures S2Eiv-v**), while soleus-innervating motor axons remained similarly unresponsive (**Figures S2Dii-iii**). Such insensitivity to BDNF stimulation is most striking when comparing transport dynamics in TA-innervating motor axons stimulated with BDNF in WT vs. SOD1^G93A^ mice (**Figure 2C**).

Collectively, these data demonstrate MN subtype-specific alterations in transport of signalling endosomes in SOD1^G93A^ mice, and that diseased TA-innervated FMNs develop a preferential insensitivity to BDNF stimulation.

### TrkB.T1 and p75^NTR^ expression is increased in SOD1^G93A^ muscles, but not at NMJs

Next, we evaluated the expression of BDNF and its receptors in TA and soleus muscles of WT and SOD1^G93A^ mice using western blot (**Figure S3A**). We found greater basal BDNF expression in TA compared to soleus, without alterations in disease (**Figure 3Ai**). There were also no discernible differences in TrkB.FL expression (**Figure 3Aii**); however, the expression of TrkB.T1 (**Figure 3Aiii**) and p75^NTR^ (**Figure 3Aiv**) was upregulated in SOD1^G93A^ muscles.

**Figure 3.**
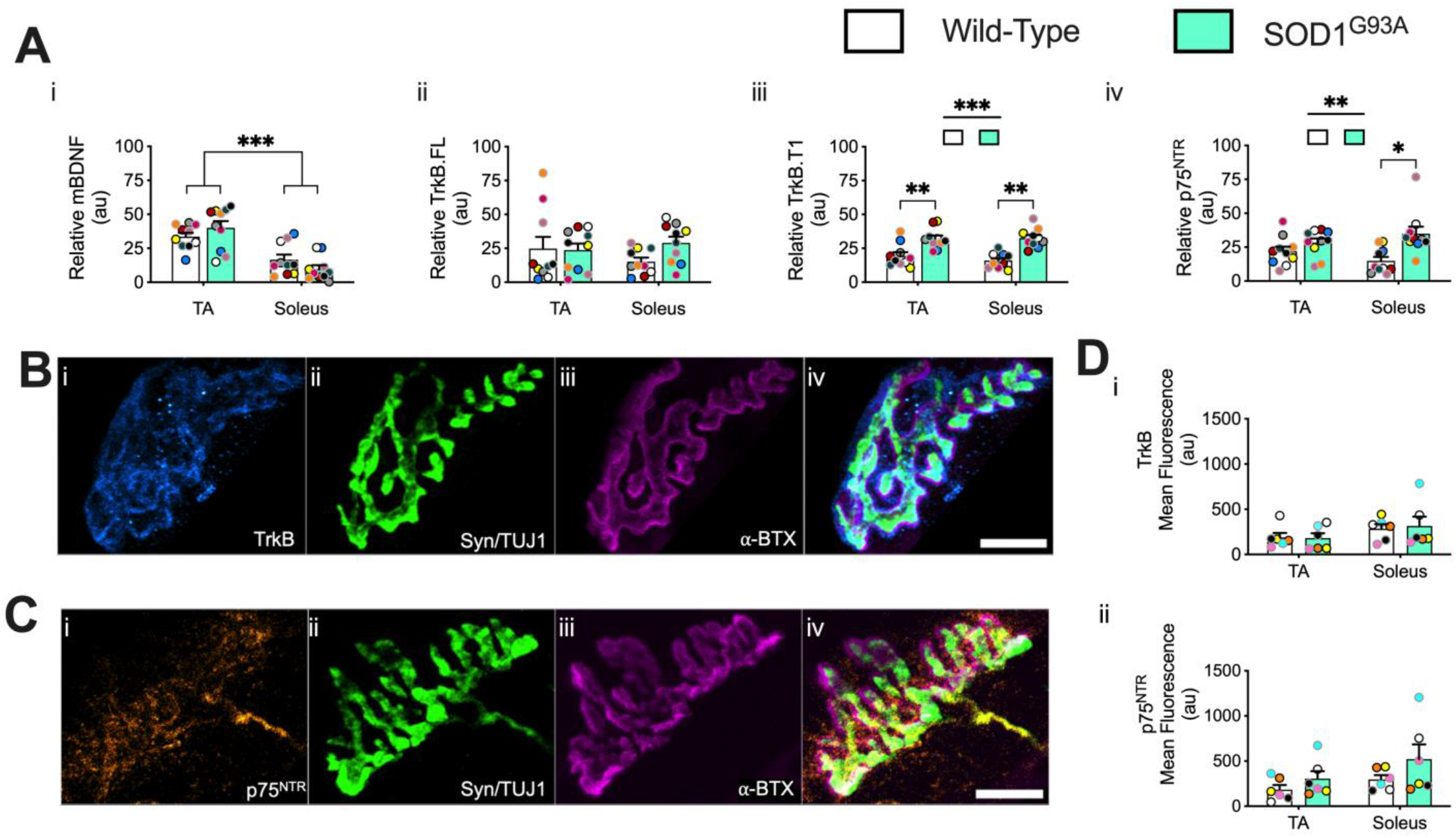
TrkB.T1 and p75^NTR^ display increased expression in SOD1^G93A^ muscles. **A)** Western blot analyses in TA and soleus muscles in WT and SOD1^G93A^ mice: *i)* mature BDNF (mBDNF) (genotype p=0.95, muscle p<0.001, interaction p=0.08), *ii)* TrkB.FL (genotype p=0.476, muscle p=0.342, interaction p=0.507), *iii)* TrkB.T1 (genotype p<0.0001, muscle p=0.239, interaction p=0.384) and *iv)* p75^NTR^ (genotype p=0.008, muscle p=0.932, interaction p=0.14) (n=10). **B)** NMJ immunostainings of TrkB and **C)** p75^NTR^ receptors at the pre- and post-synapse as identified by Syn/TUJ1 and α-BTX, respectively. **D)** Immunostaining quantifications of *i)* TrkB (genotype p=0.868, muscle p=0.109, interaction p = 0.781) and *ii)* p75^NTR^ receptors (genotype p=0.091, muscle p=0.108, interaction p=0.603) (n=6). Data were compared by two-way ANOVA and Holm-Šídák’s multiple comparisons test (A, D). *p<0.05, **p<0.01, ***p<0.001. Means ± SEM are plotted for all graphs. Black (P73) and grey (P94) circle borders indicate age-matched mice. Scale bars = 10 μm. See also **Figure S3**.

To determine if the observed changes in neurotrophin receptors were confined to the synapse, we evaluated the synaptic expression of total TrkB and p75^NTR^ at the NMJ using immunostaining. Pre-synaptic axon terminals were identified by synaptophysin (Syn) and βIII-tubulin (TUJ1), while the post-synaptic region was labelled with α-bungarotoxin (α-BTX). Applying a TrkB (**Figure 3B**) or p75^NTR^ mask (**Figure 3C**) to partially or fully innervated, but not vacant, NMJs, we found that the mean fluorescence of TrkB (**Figure 3Di; Figure S3B**) or p75^NTR^ (**Figure 3Dii; Figure S3C**) did not differ between TA or soleus NMJs in WT or SOD1^G93A^ mice. Altogether, these data reveal that while TrkB.T1 and p75^NTR^ expression is increased in SOD1^G93A^ whole muscles, this increase is not reflected at the NMJ.

### SOD1^G93A^ sciatic nerves display FMN axon diameter decreases and upregulated TrkB.T1 and p75^NTR^ expression

In agreement with previous reports that TA motor units are preferentially vulnerable in ALS mice (Nijssen et al., 2017), we found a reduction in the mean diameters in TA motor axons at P73 (**Figure 4Ai**) that persisted to P94 (**Figure 4Aii**). Suggestive of delayed pathology, LG motor axons displayed diameter reductions at P94 only (**Figure 4Aii**). Consistent with our transport data, soleus motor axons did not display altered axonal diameters in disease (**Figure 4A**).

**Figure 4.**
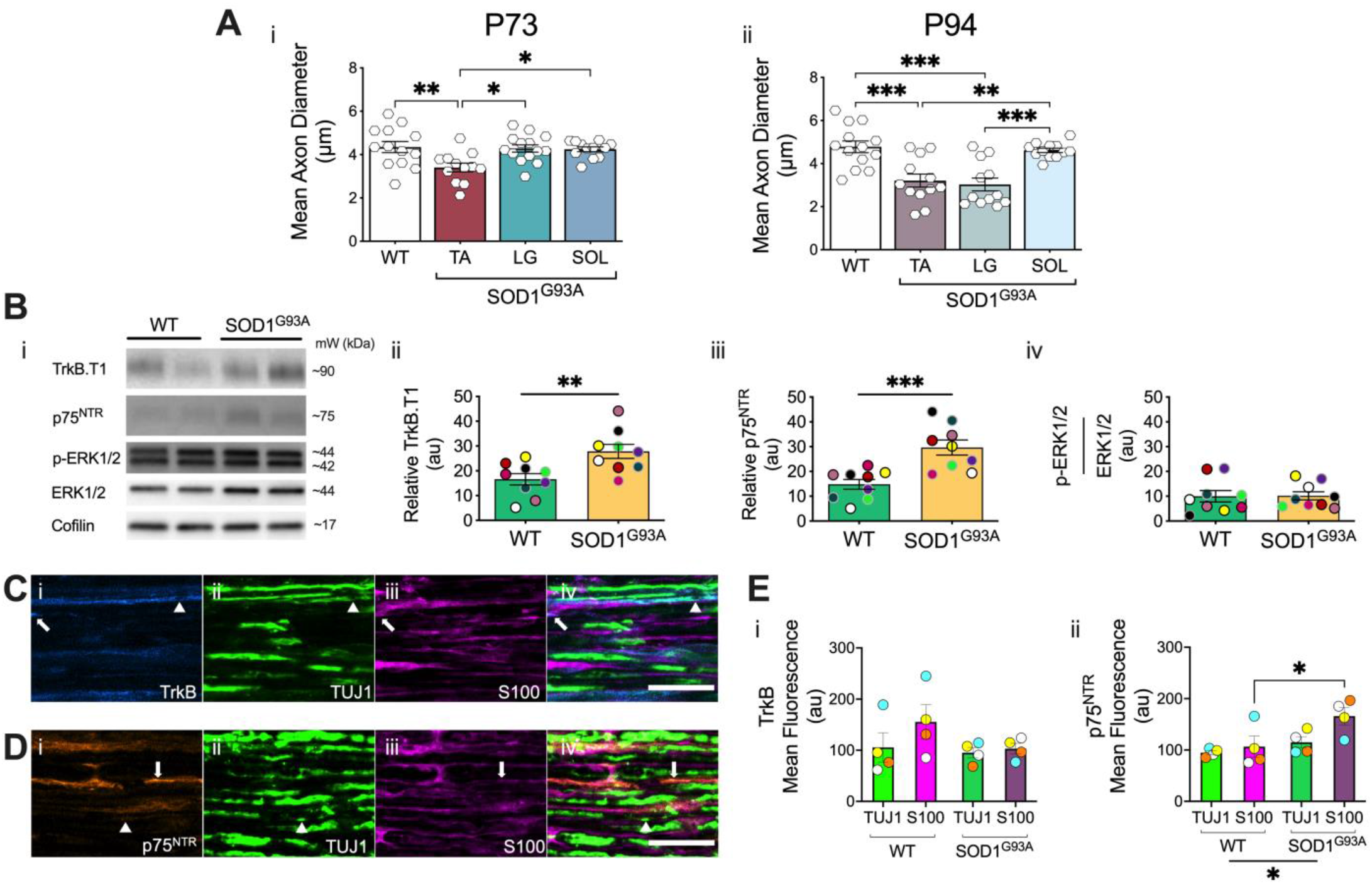
FMN axon diameters decrease whilst TrkB.T1 and p75^NTR^ increase in SOD1^G93A^ sciatic nerves. **A)** Mean diameter of H_C_T-containing motor axons innervating TA, LG, and Sol in SOD1^G93A^ and wild-type (WT) sciatic nerves at *i)* P73 (p = 0.004) and *ii)* P94 (p < 0.0001), as assessed by one-way ANOVA and Holm-Šídák’s multiple comparisons test (n=12-14). **B)** *i)* Immunoblots of TrkB.T1, p75^NTR^, p-ERK1/2, ERK1/2, and cofilin from WT and SOD1^G93A^ sciatic nerves. *NB*. TrkB.FL expression was below detection levels. Immunoblot quantifications of *ii)* TrkB.T1 (p = 0.007), *iii)* p75^NTR^ (p = 0.0008) and *iv)* p-ERK1/2-ERK1/2 ratio (p = 0.948), as compared by t-tests (n=10). Black (P73) and grey (P94) circle borders indicate age-matched mice. **C)** Immunostaining of TrkB and **D)** p75^NTR^ receptors in axons (TUJ1) and Schwann cells (S100). Arrows and arrowheads identify TrkB/p75^NTR^ expression in S100 or TUJ1 regions, respectively. Scale bar = 25 μm. **E)** Immunostaining quantifications of *i)* TrkB (genotype p = 0.204, cell type p = 0.239, interaction p = 0.384) and *ii)* p75^NTR^ (genotype p = 0.018, cell type p = 0.053, interaction p = 0.204) levels in axons or Schwann cells, as assessed by two-way ANOVA and Holm-Šídák’s multiple comparisons test (n=4). Means ± SEM are plotted for all graphs. *p<0.05, **p<0.01, ***p<0.001.

We then performed western blot analyses on whole sciatic nerves, probing for the TrkB and p75^NTR^ receptors, as well as the phosphorylation of key downstream signalling molecules, ERK1/2 and Akt (**Figure 4Bi**). Consistent with our muscle data (**Figure 3A**), TrkB.T1 (**Figure 4Bii**) and p75^NTR^ (**Figure 4Biii**) expression is increased in SOD1^G93A^ sciatic nerves, without p-ERK1/2 relative to total ERK1/2 alterations (**Figure 4Biv**). However, we were unable to detect TrkB.FL, p-AKT or AKT in our experimental conditions. To pinpoint the cellular source of the TrkB.T1 and p75^NTR^ receptors, we immunostained sciatic nerve cryosections for TrkB (**Figure 4C**) and p75^NTR^ (**Figure 4D**), using TUJ1 and S100 as markers of axons and Schwann cells, respectively. We then applied a TrkB (**Figure 4C**) or p75^NTR^ mask (**Figure 4D**) to the TUJ1 and S100 immunostained regions in WT and SOD1^G93A^ sciatic nerves. This analysis revaled no differences in TrkB content in axons or Schwann cells in WT and SOD1^G93A^ sciatic nerves (**Figure 4Ei**). In contrast, we observed increased p75^NTR^ mean fluorescence, specifically in Schwann cells (**Figure 4Eii**). Collectively, vulnerable MNs display decreases in axon diameters, and further identify non-cell autonomous, dysregulated TrkB.T1 and p75^NTR^ signalling contributions in pathology.

## Discussion

Here, we show that FMN and SMN axons have similar transport kinetics of signalling endosomes under basal conditions, with BDNF boosting axonal transport exclusively in FMNs. In SOD1^G93A^ mice, axonal transport is selectively impaired in FMN-innervated TA, which also develop an insensitivity to BDNF stimulation. Moreover, pathology increases TrkB.T1 and p75^NTR^ expression in both muscle and sciatic nerve, but not at the NMJ. Altogether, these data indicate that cell- and non-cell autonomous BDNF signalling is impaired in SOD1^G93A^ pathology, in an α-MN subtype-specific manner.

### α-MN axon subtypes display distinct transport dynamics in WT and SOD1^G93A^ mice

This is the first study to dissect axonal transport dynamics in different α-MN subtypes. In WT mice, we found no differences in speeds, but signalling endosomes paused less in TA-innervating axons. BDNF stimulation specifically enhanced axonal transport in FMNs, an effect not observed with GDNF stimulation. This supports previous reports that individual neurotrophins elicit discrete signalling in MN subtypes (Schaller et al., 2017), but is the first to account for their influence on axonal transport dynamics *in vivo*.

We have previously demonstrated that axonal transport is impaired in pre-symptomatic SOD1^G93A^ (Bilsland et al., 2010) and TDP-43^M337V^ mice (Sleigh et al., 2020b); however, compromised axonal transport is not a general by-product of neuromuscular disease as heterozygous mutant FUS (Sleigh et al., 2020b) and Kennedy’s disease (Malik et al., 2011) mice do not display *in vivo* transport deficits, despite MN loss. However, in these studies axonal transport was assessed after both TA and LG were stimulated with BDNF. As only TA axons display transport deficits in SOD1^G93A^ mice and without pausing alterations, suggest this is not due to a general impairment in the retrograde transport machinery. Interestingly, the TA-transport deficits did not worsen during disease, indicative of a pathological plateau, which was also observed in TDP-43^M337V^ mice (Sleigh et al., 2020b). However, we are unable to account for transport dynamics in denervated MNs, as only axons with internalised H_C_T could be assessed.

The precise mechanism by which FMNs selectively display impairments in retrograde axonal transport remains elusive. Mutant SOD1 protein interacts with dynein motor complexes (Zhang et al., 2007), suggesting that vulnerable FMNs may accumulate more mutant SOD1 protein, thus impinging their retrograde transport regulation. Mutant SOD1 also aberrantly interacts with the stress granule protein G3BP1 (Gal et al., 2016), thus potentially disturbing axonal maintenance functions, (e.g., stress granule dynamics, altered RNA localisation/availability) (Sahoo et al., 2018). Axonal transport deficits also impact local translation, as Rab7-containing organelles, which include signalling endosomes, are sites for mitochondrial-associated local mRNA translation (Cioni et al., 2019). Whether these pathological phenomena occur specifically in vulnerable FMNs, but not in resistant SMNs, or whether other organelles also display transport deficits specifically in vulnerable FMNs, remains to be determined.

### Dysregulated TrkB.T1 and p75^NTR^ expression in ALS mice

Dynamic NMJ remodelling precedes motor unit loss in ALS mice (Tremblay et al., 2017; Martineau et al., 2018), however it is currently not known whether neuromuscular BDNF signalling in fast vs. slow muscles is altered in disease. Remarkably, we report that BDNF signalling in SOD1^G93A^ mice is dysregulated in embryonic neurons, and adult MNs are insensitivity to BDNF stimulation, with TA axons displaying ∼38% reduction in transport speeds in early symptomatic (P73) SOD1^G93A^ mice. As physiological BDNF and TrkB levels fluctuate (e.g., increase with exercise (Gómez-Pinilla et al., 2002; Ferraiuolo et al., 2009; Hurtado et al., 2017)), persistent BDNF insensitivity can have severe consequences for MN homeostasis, impacting translation and signalling events in axon terminals, along the axon and within MN soma (Santos et al., 2010; Hurtado et al., 2017; Delezie et al., 2019; Bronfman and Moya-Alvarado, 2020; Kim and Jung, 2020).

We initially hypothesised that this BDNF insensitivity might be due to: 1) reduced muscle BDNF expression; 2) disproportionate TrkB and p75^NTR^ levels; 3) imbalanced TrkB.FL and TrkB.T1 ratios; or 4) a combination of the above. However, the only observed pathological alteration identified in our study was an increase in TrkB.T1 and p75^NTR^ expression in SOD1^G93A^ sciatic nerves and muscles, suggestive of muscle and nerve TrkB.T1 and p75^NTR^ contributions to pathology. However, a caveat to our TrkB staining approach is that the commercially available TrkB antibodies bind to the extracellular domain of this receptor, and thus cannot distinguish TrkB.FL from TrkB.T1. Our experimental conundrum is that while western blotting allows us to evaluate the expression of the TrkB isoforms, it lacks MN subtype specificity; conversely, our immunostaining experiments enable MN subtype specificity, but lack TrkB isoform differentiation. Hence, dissecting the endogenous expression of TrkB isoforms in FMNs and SMNs is currently not possible.

MN-specific TrkB deletion reduces mutant SOD1 protein inclusions and axonal degeneration (Zhai et al., 2011). In addition, α-MNs upregulate p75^NTR^ and apoptotic markers in symptomatic SOD1^G93A^ mice (Smith et al., 2015), and ubiquitous TrkB.T1- or astrocyte-specific TrkB.T1-deletion delays MN death in SOD1^G93A^ mice (Yanpallewar et al., 2012; 2021), suggesting that modulating these classical non-pro-survival receptors might have therapeutic benefits. Indeed, harnessing the pro-survival activity of p75^NTR^ prevents MN death and extends the lifespan of SOD1^G93A^ mice by partially rescuing reduced p-TrkB, p-Akt, p-ERK and p-CREB expression in SOD1^G93A^ spinal cords (Matusica et al., 2016).

Collectively, the BDNF signalling axis appears essential for maintenance and homeostatic regulation of FMNs, which is selectively impaired in SOD1^G93A^ pathology with cell- and non-cell autonomous alterations.

### Therapeutic Relevance

In our study, we reveal an impairment in BDNF signalling that occurs specifically in vulnerable FMNs, thus laying the foundations for future therapeutic interventions for ALS. Moreover, therapeutically targeting axonal transport appears to be a viable strategy to promote MN health in ALS. Indeed, acute modification of growth factor receptor activation (e.g., IGF1R) or kinase inhibition (e.g., p38 MAPK) rescued axonal transport deficits *in vitro* and *in vivo* in SOD1^G93A^ mice (Gibbs et al., 2018; Fellows et al., 2020), and an isoform-specific p38 MAPK inhibitor is currently in clinical trials for dementia with Lewy bodies (NCT02423200). Alternatively, gene therapy offers a long-term strategy to enhance pro-survival signalling and/or reduce gene mutation toxicity (Tosolini and Sleigh, 2017). In particular, simultaneous overexpression of BDNF and TrkB produces the best enhancement of optic nerve anterograde transport in glaucoma and tauopathy models (Khatib et al., 2021), and may prove an exciting new therapeutic strategy for vulnerable MNs.

## Acknowledgements

This work is dedicated to the memory of Paul Victor Tosolini. We thank James Dick and the personnel of the Denny Brown Laboratories for assistance in maintaining the mouse colonies (Queen Square Institute of Neurology, University College London), and Nicol Birsa, Jose Norberto Sagullo Vargas and David Villarroel-Campos (Queen Square Institute of Neurology, University College London) for critical reading of the manuscript. This work was supported by a Junior Non-Clinical Fellowship from the Motor Neuron Disease Association (Tosolini/Oct20/973-799) (APT); the Medical Research Council Career Development Award (MR/S006990/1) (JNS); the Sir Henry Wellcome fellowship 103191/A/13/Z (JNS); Human Frontier Science Program long-term fellowship LT000220/2017-L (SS); Medical Research Council Studentship (ERR); Horserace Betting Levy Board and the Mellow foundation provided salary support (SDB); Wellcome Senior Investigator Awards (107116/Z/15/Z and 223022/Z/21/Z) (GS), and a UK Dementia Research Institute Foundation award (UKDRI-1005) (GS).

## Author Contributions

Conceptualisation: APT and GS. Investigation APT, JNS, SS, ERR and SC. Writing and figure production: APT and GS, with input from all authors. Funding acquisition: APT and GS. All authors approved this submission.

## Materials and Methods

### Animals

Mouse experiments were performed under license from the United Kingdom Home Office in accordance with the Animals (Scientific Procedures) Act (1986) and approved by the UCL Queen Square Institute of Neurology Ethics Committee. Mice were housed in individually-ventilated cages in a controlled temperature/humidity environment and maintained on a 12 h light/dark cycle with access to food and water provided *ad libitum*. Transgenic mice carrying the mutant SOD1^G93A^ transgene (TgN[SOD1-G93A]1Gur) gene (Gurney et al., 1994) were obtained from the Jackson Laboratories. Colonies were maintained by breeding male heterozygous carriers with female (C57BL/6 x SJL) F1 hybrids. Mice were genotyped for the human SOD1 transgene using DNA extracted from ear notches and primers as previously described (Bilsland et al., 2010; Gibbs et al., 2018; Fellows et al., 2020). For all experiments, only female hemizygous transgenic mice carrying the human SOD1^G93A^ transgene (referred hereafter as SOD1^G93A^) or wild-type (WT) littermates were used. For axonal transport, female WT mice had a mean age of 85.13 ± 14.97 days, which is comparable across both mutant timepoints, as age does not influence *in vivo* signalling endosome dynamics (Sleigh et al., 2020b). Female SOD1^G93A^ postnatal day 73 (P73) mice had a mean age of 72.74 ± 0.67 and postnatal day 94 (P94) mice had a mean age of 93.70 ± 0.46.

### In vivo axonal transport

Signalling endosomes were visualised *in vivo* by injecting the fluorescent atoxic binding fragment of tetanus neurotoxin (H_C_T-555) as previously described (Sleigh et al., 2020a). Briefly, H_C_T (residues 875-1315) fused to an improved cysteine-rich region was expressed in bacteria as a glutathione-S-transferase fusion protein (Restani et al., 2012), and subsequently labelled with AlexaFluor555 C_2_ maleimide (Thermo Fisher Scientific, A-20346). 5-7.5 μg of H_C_T-555 alone, or in combination with 25 ng of human recombinant BDNF (Peprotech, 450-02) or 25 ng of human recombinant GDNF (Peprotech, 450-10) (pre-mixed with phosphate buffered saline) were injected in single muscles. *In vivo* axonal transport experiments were performed as previously described (Sleigh et al., 2020a). Briefly, after anaesthesia was initiated and maintained using isoflurane, the fur on the ventral and/or dorsal lower leg was shaved off, and mice were placed on a heat-pad for the entire duration of the surgery. A small incision was made using iris spring scissors on the ventral surface below the patella for tibialis anterior (TA), or on the lateral aspect of the dorsal surface below the popliteal fossa for lateral head of gastrocnemius (LG). Injections were performed as a single injection targeting the motor end plate region (Mohan et al., 2014) in a volume of ∼3.5 μl using a 701N Hamilton® syringe (Merck, 20779) for TA and LG. For soleus injections, a vertical incision was made using iris spring scissors on the skin covering the lateral surface of lower hindlimb between the patella and tarsus to expose the underlying musculature. Subsequent vertical incisions were carefully made laterally along the connective tissue between LG and TA, and the deeper soleus muscle was exposed using forceps. 1 μl injections were performed into soleus using pulled graduated, glass micropipettes (Drummond Scientific, 5-000-1001-X10), as previously described (Mohan et al., 2015). The overlying skin was then sutured, and mice were monitored for up to 1 h. 4-8 h later, mice were re-anaesthetised with isoflurane, and the skin covering the entire lateral surface of the injected hindlimb was removed, along with the biceps femoris muscle to expose the underlying sciatic nerve. The connective tissue underneath the sciatic nerve was disrupted using curved forceps to enable the placement of a small piece of parafilm aiding the subsequent imaging. The anaesthetised mouse was then transferred to an inverted LSM780 laser scanning microscope (Zeiss) enclosed within an environmental chamber pre-warmed and maintained at 37°C. Using a 40x, 1.3 NA DIC Plan-Apochromat oil-immersion objective (Zeiss), axons containing retrogradely motile H_C_T-555-positive signalling endosomes were imaged every 0.3-0.4 s using a 80x digital zoom (1024 × 1024, <1% laser power) (**Figure 1A**,); movies of three to five axons per animal were acquired. All imaging was concluded within 1 h of reinitiating anaesthesia.

### In vivo axonal transport analysis

Confocal “.czi” images were opened in FIJI/ImageJ (http://rsb.info.nih.gov/ij/), converted to “.tiff” and transport dynamics were then assessed semi-automatically (i.e., automated spot detection and manual linking (see Video 1) using the TrackMate plugin (Tinevez et al., 2017). Kymographs (**Figure S2A**) were generated using ImageJ but were not used to assess axonal transport dynamics. Only thicker axons that displayed the stereotypical pattern of movement of a MN, as previously determined using ChAT.eGFP mice (Sleigh et al., 2020b), were selected for tracking. Individual endosomes were tracked if there was movement in 10-100 consecutive frames, including non-terminal pauses (defined by no movement in ≤10 consecutive frames), but excluding terminal pauses (defined by no movement in ≥10 frames). Inclusion criteria of ≥1,000 endosomal frame-to-frame movements from a total of 15-40 endosomes across ≥3 axon bundles were analysed per individual animal (i.e., all those criteria were met for n=1). The breakdown of each experimental group can be found in Table S1. Relative frequency graphs were generated to represent the relative frame-to-frame movements of all signalling endosomes per animal. The mean endosome speed was determined by averaging the mean speeds of all tracked endosomes from every axon assessed per animal. The fastest average endosome speed of a single endosome per animal was considered as the maximum speed. A pause was defined by an endosome that moved <0.1 μm in consecutive frames, and the time paused (%) is determined by the number of pauses divided by the total number of frame-to-frame movements assessed per animal.

### In vitro axonal transport

Mixed ventral horn cultures were prepared as previously described (Gibbs et al., 2018; Fellows et al., 2020). Briefly, ventral horns from E11.5-13.5 SOD1^G93A^ and WT mice were dissociated, centrifuged at 380 g for 5 min, seeded into two-chambered microfluidic devices (**Figure S2Bi**) (Fellows et al., 2020), and maintained in motor neuron media (Neurobasal (Gibco) with 2% B27 (Gibco), 2% heat-inactivated horse serum, 1% Glutamax (Invitrogen), 24.8 μM β-mercaptoethanol, 10 ng/ml ciliary neurotrophic factor (Peprotech, 450-13), 0.1 ng/ml GDNF (Peprotech, 450-10), 1 ng/ml BDNF (Peprotech, 450-02) and 1x penicillin streptomycin (Thermo Fisher; 15140122)) at 37°C and 5% CO_2_. After 6 days *in vitro*, 30 nM H_C_T-555 and +/-50 ng/ml of BDNF was added to existing media for 45 min, then all media was replaced with fresh MN media containing 20 mM HEPES-NaOH (pH 7.4) +/-50 ng/ml of BDNF for time-lapse microscopy. Live imaging was performed on the same inverted LSM780 laser scanning microscope (Zeiss) enclosed within an environmental chamber pre-warmed and maintained at 37°C and imaged using a 40x, 1.3 NA DIC Plan-Apochromat oil-immersion objective (Zeiss). Videos were taken at 500 ms frame rate for >2.5 min. Videos were manually tracked using TrackMate (Tinevez et al., 2017) to determine endosome track dynamics (see **Figure 2B** legends for precise details).

### Axon diameters

The axon diameters were measured following protocols established in ChAT.eGFP mice (Sleigh et al., 2020b), using the same videos in the axonal transport analyses. Briefly, axon diameters were assessed by measuring the upper and lower positions of moving H_C_T-555 signalling endosomes from consecutive frames in unprocessed (i.e., not dissected, fixed, or sectioned), anatomically connected individual axons. A minimum of 10 positions were averaged for a single axon, and the mean axon diameters per animal was determined by averaging all axons from that animal (n ≥3 axons per animal). Similar to the axonal transport experiments, this assessment is reliant upon intact NMJs, which can internalise H_C_T-555; hence we cannot extrapolate diameters from denervated axons.

### Muscle BDNF, TrkB and p75^NTR^ expression

P73 (n=5) and P94 (n=5) WT and SOD1^G93A^ mice were culled, and fresh TA and soleus muscles were immediately dissected, snap frozen in liquid nitrogen and stored at -80°C. Protein extraction from frozen muscles was achieved by mechanically disrupting the tissue using a scalpel, followed by immersion in RIPA buffer (50 mM Tris-HCl pH 7.5, 1 mM EDTA, 2 mM EGTA, 150 mM NaCl, 1% NP40, 0.5% sodium deoxycholate, 0.1% SDS) containing Halt protease and phosphatase inhibitor cocktail (10% weight/volume) (100x, Fisher, 78442) for 15 min on ice, and then homogenised on ice using an electrical homogeniser. Lysates were incubated at 4°C with mild agitation for 2 h, after which they were centrifuged at 21,000g for 30 min at 4°C. 20 μl of each supernatant (containing ∼60 μg of protein) was treated with 6.5% trichloroacetic acid and acetone. The protein pellet was resuspended in 1x Laemmli buffer and loaded on Bis-Tris polyacrylamide gels. Western blotting was then performed using standard protocols. Primary antibodies against BDNF (Alomone, ANT-010, 1:1000), TrkB (Millipore, 07-225, 1:1000), and p75^NTR^ (Biolegend, 839701, 1:1000) were used to quantify protein levels. To avoid muscle-specific differences in the levels of housekeeping gene(s) (Wyckelsma et al., 2016) mBDNF, TrkB.FL, TrkB.T1 and p75^NTR^ bands were first standardised to the total protein using Coomassie staining on the blot, and then normalised by the sum of all data points in a replicate (Degasperi et al., 2014). P73 and P94 WT and SOD1^G93A^ data points were combined as there were no timepoint specific differences (data not shown).

### Muscle immunohistochemistry

P73 (n=3) and P94 (n=3) WT and SOD1^G93A^ mice were culled, and TA and soleus muscles were immediately dissected and post-fixed in 4% paraformaldehyde (PFA) for 15-60 min. Muscle fibres were teased apart in bundles of 1-10 fibres and stained with α-bungarotoxin (Thermo Fisher Scientific, B13423, 1:500) for 1 h. Fibres were then permeabilized with 2% Triton X-100 in PBS for 90 min, then immersed in a blocking solution containing 4% bovine serum albumin and 1% Triton X-100 in PBS for 30 min at room temperature. Primary antibodies staining for TUJ1 (Synaptic systems, 302306, synaptophysin, 1:50), TrkB (Millipore, 07-225, 1:50) and p75^NTR^ (Promega, G3231, 1:50) immersed in blocking solution were added to the teased muscle fibres for ∼3 d at 4°C with mild agitation, and then washed in PBS at room temperature. Secondary antibodies in PBS were then applied to fibres for ∼1 h at room temperature, followed by multiple washes in PBS and then finally mounted on SuperFrost Plus slides (VWR, 631-0108) using Mowiol. Slides were dried and imaged with a LSM780 laser scanning microscope using a 63x Plan-Apochromat oil immersion objective (Zeiss). A minimum of 25 NMJs were imaged per condition, comprised of fully or partially innervated, but not denervated, NMJs. Mean fluorescence was measured using FIJI/ImageJ by applying a TrkB or p75^NTR^ mask to the overlapped BTX-synaptophysin/TUJ1 NMJ region, and the mean fluorescence per animal was assessed by averaging all individual data points. For the TrkB analysis, we assessed the WT TA (n=6, NMJs = 182), WT soleus (n=6, NMJs = 223), SOD1^G93A^ TA (n=6, NMJs = 180) and SOD1^G93A^ soleus (n=6, NMJs = 191) muscles. For the p75^NTR^ analysis, we assessed the WT TA (n=6, NMJs = 176), WT soleus (n=6, NMJs = 224), SOD1^G93A^ TA (n=6, NMJs = 183) and SOD1^G93A^ soleus (n=6, NMJs = 198) muscles. P73 and P94 WT and SOD1^G93A^ data points were combined as there were no time-point specific differences (data not shown).

### Sciatic nerve expression

P73 (n=5) and P94 (n=5) WT and SOD1^G93A^ mice were culled, and sciatic nerves were immediately dissected, snap frozen in liquid nitrogen and stored at -80°C. Thawed sciatic nerves were then immersed in NP-40 lysis buffer (150 mM NaCl, 1% NP-40, 50 mM Tris-HCl, pH 8.0) Halt protease and phosphatase inhibitor cocktail (10% weight/volume) (100x, Fisher, 78442). Lysates underwent mechanical disruption using a plastic pestle before being left on ice for 0.5 h and then centrifuged for 20 min at 10,000*g*. The protein pellet was re-suspended and denatured in 4x Laemmli buffer, loaded on Bis-Tris polyacrylamide gels prior to western blotting. Primary antibodies against TrkB (Millipore, 07-225, 1:1000), p75^NTR^ (Biolegend, 839701, 1:2000), ERK1/2 (CST, 9102, 1:1000), p-ERK1/2 (CST, 9101, 1:1000), AKT (CST, 9272, 1:1000), p-AKT (CST, 9275, 1:1000) and Cofilin (Cytoskeleton, ACFL02, 1:500) were used to quantify protein levels. *N*.*B*. The expression of TrkB.FL, AKT and p-AKT were below detection levels. All bands were first standardised to Cofilin, and then normalised by the sum of all data points in a replicate (Degasperi et al., 2014). P73 and P94 WT and SOD1^G93A^ data points were combined as there were no time-point specific differences (data not shown).

### Sciatic nerve IHC

P73 (n=4) WT and SOD1^G93A^ mice were culled, and sciatic nerves were immediately dissected, post-fixed in a 4% PFA solution in PBS overnight at 4°C, cryopreserved in a 30% sucrose solution in PBS for 2 d at 4°C, and finally frozen in OCT (Agar Scientific, AGR1180). 30 μm cryosections of sciatic nerves were directly mounted on SuperFrost Plus slides (VWR, 631-0108), and a hydrophobic barrier pen (Vector Laboratories, H-4001) was then applied to the slides surrounding the sectioned tissue. PBS rehydrated tissue was then blocked using 10% normal horse serum in 0.2% Triton X-100 in PBS for ∼1 h and then primary antibodies specific for S100 (Merck, S2532, 1:200) TUJ1 (Synaptic systems, 302306, synaptophysin, 1:500), TrkB (Millipore, 07-225, 1:250) and p75^NTR^ (Promega, G3231, 1:500) were applied overnight at room temperature. After multiple PBS washes, the secondary antibodies in PBS were applied for 2-3 h, followed by multiple washes and then mounted with Mowiol. Slides were dried and imaged with a LSM780 laser scanning microscope using a 63x Plan-Apochromat oil immersion objective. A minimum of 6 sciatic nerve sections were imaged per condition. Mean fluorescence was measured by applying a TrkB or p75^NTR^ mask to the TUJ1 and S100 regions, and the mean fluorescence per animal was assessed by averaging all the individual data points.

## Statistical analysis

GraphPad Prism 9 (GraphPad Software, San Diego, CA, USA) was used for statistical analyses. Normal distribution was first ascertained by the D’Agostino and Pearson omnibus normality test, and parametric data were statistically assessed using unpaired, two-tail t-tests, one-way or two-way analyses of variance (ANOVA) with Holm-Sidaks multiple comparison tests. Non-normally distributed data were analysed by a two-tailed Mann-Whitney *U* test or Kruskal-Wallis test with Dunn’s multiple comparisons test.

