## Supplementary Information for "BDNF-dependent modulation of axonal transport is selectively impaired in ALS"

### Supplementary Figures

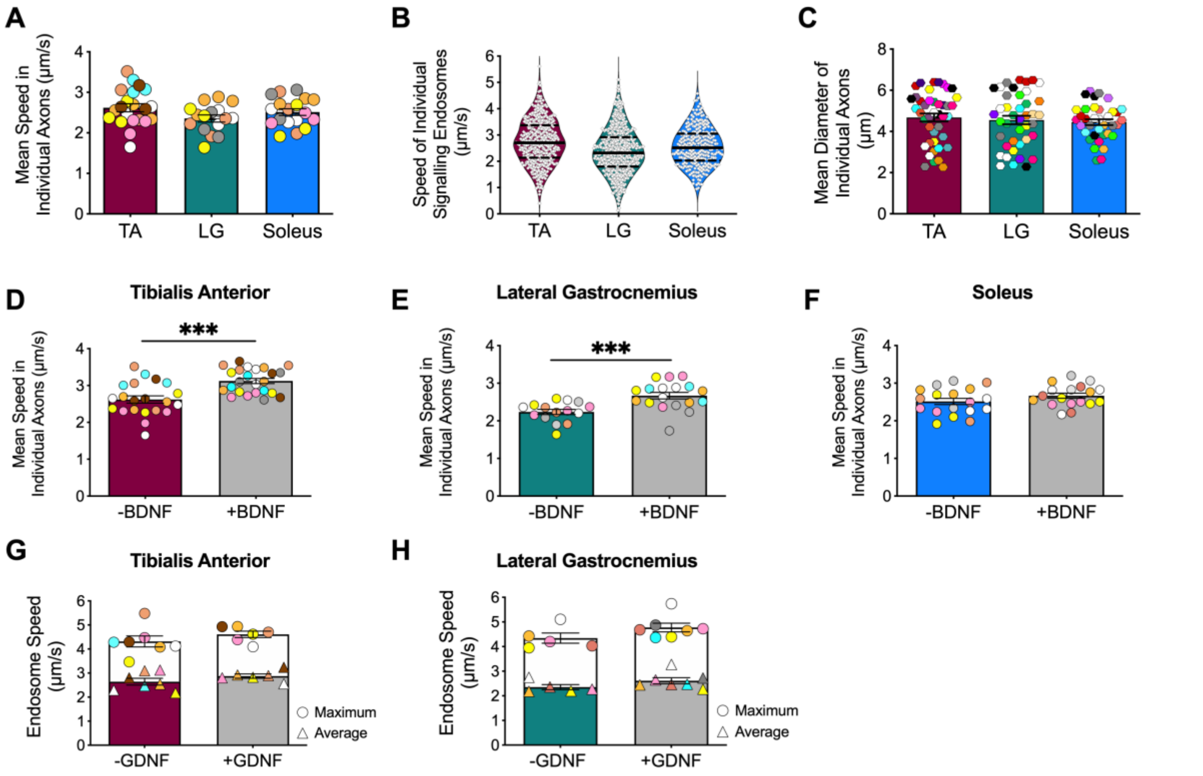


**Supplementary Figure 1. Wild-type retrograde transport dynamics of signalling endosomes in single axons. A)** Mean speeds of $H_C$T-555-positive signalling endosomes in individual motor neuron axons innervating tibialis anterior (TA), lateral gastrocnemius (LG) and soleus ($p$ = 0.135, one-way ANOVA, n=16-21). The colour coding is consistent with **Figures 1C-D**. **B)** Speed of individual $H_C$T-555-positive signalling endosomes (white diamonds) in motor axons innervating TA, LG, and soleus. Black line represents the median and the dashed line represents the upper and lower quartiles. **C)** Mean diameters of individual motor axons innervating TA, LG and soleus that contain $H_C$T-555-positive signalling endosomes ($p$=0.695, one-way ANOVA, n=36-42). The colour coding is consistent with **Figure 1E**. **D)** Mean axonal endosome speeds upon intramuscular BDNF stimulation in individual motor neurons innervating TA ($p$=0.0001), **E)** LG ($p$=0.007), and **F)** soleus ($p$=0.161), as assessed by a two-tailed unpaired t-test (n=15-24). The colour coding is consistent with **Figures 1F-K**. **G)** Mean (triangles) and maximum (circles) axonal endosome speeds upon intramuscular GDNF stimulation in TA ($p$=0.181) and **H)** LG wild-type motor neurons ($p$=0.149), as assessed by a two-tailed unpaired Mann-Whitney U test (n=6-7). Means ±SEM are plotted for all graphs. Linked to **Figure 1**.

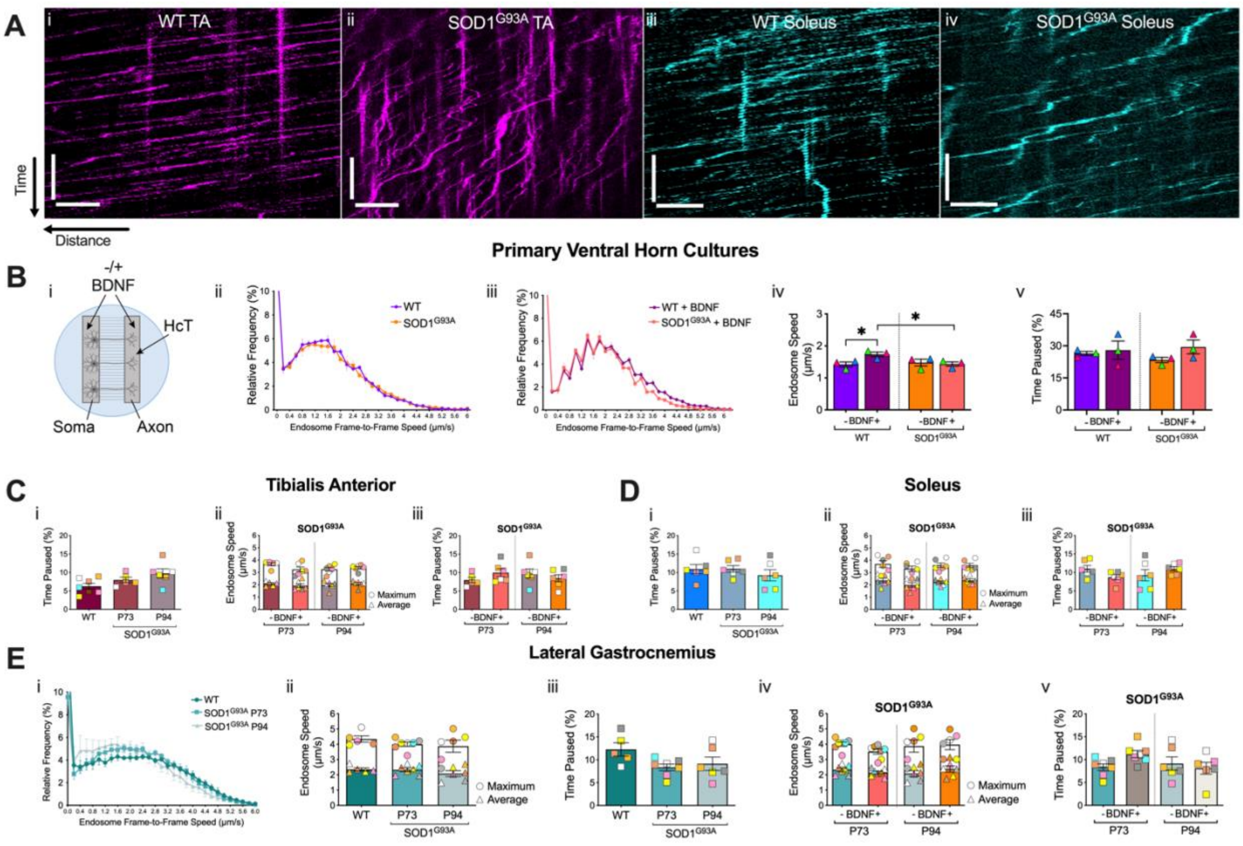


**Supplementary Figure 2. Axonal transport dynamics of signalling endosomes in SOD1$^{G93A}$ mice. A)** Representative kymographs of retrograde transport of HCT-555-positive signalling endosomes in motor axons innervating the tibialis anterior (TA) muscle in *i)* wild-type (WT) and *ii)* SOD1$^{G93A}$ mice, and motor axons innervating the soleus muscle in *iii)* wild-type and *iv)* SOD1$^{G93A}$ mice. Retrogradely moving HCT-555-positive signalling endosomes are represented by right-to-left movements and paused HCT-555-positive signalling endosomes are represented by vertical lines. Time (y-axis) scale bars = 30 s, distance (x-axis) scale bars = 10 µm. **B)** *i)* Schematic representation of WT and SOD1$^{G93A}$ primary ventral horn cultures plated in microfluidic chambers (MFCs) with or without 50 ng/µl of BDNF added to both somatic and axonal compartments. *ii*) Speed distribution curves of WT and SOD1$^{G93A}$ primary ventral horn cultures grown in MFCs. WT points represent data from 24 axons, 103 endosomes and 12,650 single endosomal movements; SOD1$^{G93A}$ points represent data from 22 axons, 119 endosomes and 11,769 single endosomal movements (n=3 biological replicates). *iii)* Speed distribution curves, *iv)* mean endosome speeds (WT vs. SOD1$^{G93A}$: p=0.068; WT -/+BDNF: p=0.041; SOD1$^{G93A}$ -/+BDNF: p=0.735; SOD1$^{G93A}$ -/+BDNF: p=0.037) and *v)* relative percentage of time signalling endosomes paused (WT vs. SOD1$^{G93A}$: p=0.122; WT -/+BDNF: p=0.745; SOD1$^{G93A}$ -/+BDNF: p=0.154; SOD1$^{G93A}$ -/+BDNF: p=0.791) of WT and SOD1$^{G93A}$ primary ventral horn cultures grown in MFCs with the addition of 50 ng/µl of BDNF. Statistical analyses were performed using an unpaired, two-tailed t-test (n=3 biological replicates,). WT points +BDNF represent data from 289 endosomes in 22 axons for a total of 21,747 single endosomal movements; SOD1$^{G93A}$ points +BDNF represent data from 238 endosomes in 21 axons for a total of 20,007 single endosomal movements. **C)** *In vivo* axonal transport dynamics in SOD1$^{G93A}$ mice of TA-innervating axons and **D)** soleus-innervating axons, displaying the *i)* relative percentage of time signalling endosomes paused (TA: p=0.061; Sol: p=0.562), *ii)* mean and maximum endosomal speeds upon intramuscular BDNF stimulation (TA: p=0.464; Sol: p=0.48) and *iii)* relative percentage of time signalling endosomes paused after BDNF stimulation (TA: p=0.521; Sol: p=0.213). **E)** Axonal transport dynamics in motor axons innervating lateral gastrocnemius at two stages of SOD1$^{G93A}$ disease (P73 and P94) compared to WT mice, displaying the *i)* speed distribution curves, *ii)* mean and maximum speeds (p=0.492), *iii)* relative percentage of time signalling endosomes paused (p=0.128), *iv)* mean and maximum endosomal speeds after BDNF stimulation (p=368) and *v)* relative percentage of time signalling endosomes paused after BDNF stimulation (p=0.136). Statistics were assessed by a one-way ANOVA followed by a Kruskal-Wallis multiple comparisons test. n=5-7. Linked to **Figure 1** and **Figure 2**.

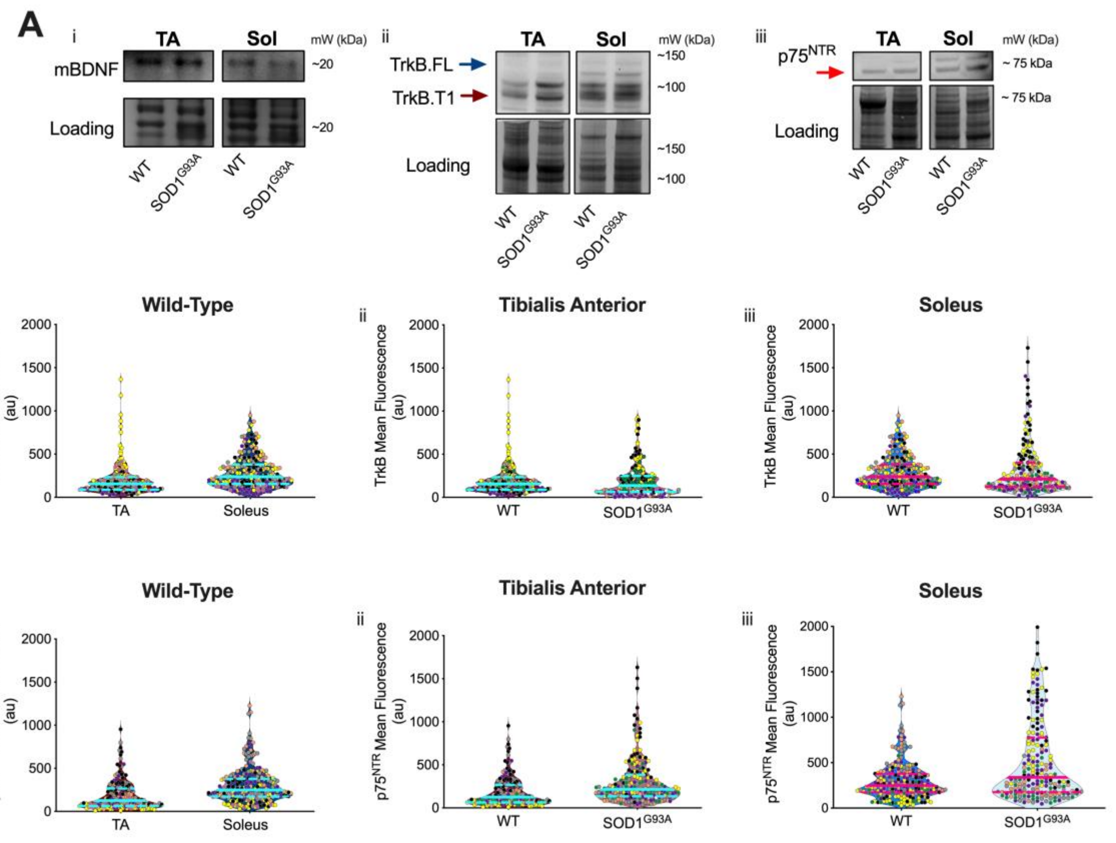


**Supplementary Figure 3. mBDNF, TrkB and p75$^{NTR}$ receptor expression in muscles and at the NMJ.**

**A)** Representative immunoblots for *i)* mature BDNF (mBDNF), *ii)* TrkB.FL and TrkB.T1, and *iii)* p75$^{NTR}$, comparing wild-type (WT) and SOD1$^{G93A}$ tibialis anterior (TA) and soleus (Sol) muscles. Linked with **Figure 3A**). **B)** *i)* Individual data points of neuromuscular junction (NMJ) TrkB mean fluorescence WT in TA (182 NMJs) and Sol (223 NMJs) muscles (n=6). Corresponds with **Figure 3Di**. *ii)* Individual data points of NMJ TrkB mean fluorescence in TA muscles comparing WT (182 NMJs) and SOD1$^{G93A}$ (180 NMJs) mice (n=6). Corresponds with **Figure 3Di**. *iii)* Individual data points of NMJ TrkB mean fluorescence in Sol comparing WT (223 NMJs) and SOD1$^{G93A}$ (191 NMJs) mice (n=6). Corresponds with **Figure 3Di**. **C)** *i)* Individual data points of NMJ p75$^{NTR}$ mean fluorescence WT TA (176 NMJs) and Sol (224 NMJs) muscles (n=6). Corresponds with **Figure 3Dii.** *ii)* Individual data points of NMJ p75$^{NTR}$ mean fluorescence in TA muscles comparing WT (176 NMJs) and SOD1$^{G93A}$ (183 NMJs) mice (n=6). Corresponds with **Figure 3Dii**. *iii)* Individual data points of NMJ p75$^{NTR}$ mean fluorescence in Sol comparing WT (224 NMJs) and SOD1$^{G93A}$ (198 NMJs) mice (n=6). Corresponds with **Figure 3Dii**. For all plots, colour coding of individual data points represents an individual animal, and match their corresponding graph in **Figures 3Di-ii**. The cyan/pink line represents the mean, and the dashed cyan/pink lines represent the upper and lower quartiles. Black (P73) and grey (P94) circles indicate age-matched mice.

| Muscle | Genotype | Number of Animals | Number of Axons | Number of signalling endosomes | Frame-to-Frames Assessed |
|---|---|---|---|---|---|
| TA | Wild type | 7 | 21 | 469 | 19,405 |
| LG | Wild type | 5 | 16 | 609 | 25,402 |
| Soleus | Wild type | 6 | 18 | 292 | 18,737 |
| TA + BDNF | Wild type | 7 | 21 | 621 | 24,543 |
| LG + BDNF | Wild type | 6 | 19 | 581 | 21,978 |
| Soleus + BDNF | Wild type | 6 | 18 | 309 | 18,932 |
| TA + GDNF | Wild type | 6 | 18 | 530 | 18,445 |
| LG + GDNF | Wild type | 7 | 29 | 947 | 30,660 |
| TA | $SOD1^{G93A}$ P73 | 5 | 15 | 269 | 15,508 |
| TA | $SOD1^{G93A}$ P94 | 6 | 18 | 309 | 18,880 |
| LG | $SOD1^{G93A}$ P73 | 7 | 21 | 430 | 21,517 |
| LG | $SOD1^{G93A}$ P94 | 6 | 18 | 408 | 18,241 |
| Soleus | $SOD1^{G93A}$ P73 | 6 | 18 | 246 | 18,416 |
| Soleus | $SOD1^{G93A}$ P94 | 6 | 18 | 238 | 18,574 |
| TA + BDNF | $SOD1^{G93A}$ P73 | 6 | 18 | 326 | 18,779 |
| TA + BDNF | $SOD1^{G93A}$ P94 | 6 | 18 | 338 | 19,009 |
| LG + BDNF | $SOD1^{G93A}$ P73 | 7 | 21 | 371 | 21,929 |
| LG + BDNF | $SOD1^{G93A}$ P94 | 6 | 18 | 394 | 18,540 |
| Soleus + BDNF | $SOD1^{G93A}$ P73 | 6 | 18 | 241 | 18,484 |
| Soleus + BDNF | $SOD1^{G93A}$ P94 | 6 | 18 | 254 | 18,493 |

**Supplementary Table 1.** Number of animals, axons, cargoes and frame-to-frame movements assessed for each *in vivo* axonal transport experimental group.
